# cGMP dynamics underlie thermosensation in *C. elegans*

**DOI:** 10.1101/764571

**Authors:** Ichiro Aoki, Makoto Shiota, Shunji Nakano, Ikue Mori

**Author notes:** Equal contribution (Ichiro Aoki, Makoto Shiota). Buchmann Institute for Molecular Life Sciences, Goethe University, Max-von-Laue-Strasse 15, D-60438 Frankfurt, Germany (Ichiro Aoki).

## Abstract

Animals sense ambient temperature so that they can adjust their behavior to the environment; they avoid noxious heat and coldness and stay within a survivable temperature range. *C. elegans* can sense temperature, memorize past cultivation temperature and navigate towards preferable temperature, for which a thermosensory neuron, AFD, is essential. AFD responds to temperature increase from the past cultivation temperature by increasing intracellular Ca2+ level. We aimed to reveal how AFD encodes and memorizes the information of temperature. Although cGMP synthesis is crucial for thermosensation by AFD, whether and how cGMP level temporally fluctuates in AFD remained elusive. We therefore monitored cGMP level in AFD and found that cGMP dynamically responded to temperature change in a manner dependent on past cultivation temperature. Given that cGMP dynamics is supposed to be upstream of Ca2+ dynamics, our results suggest that AFD’s memory is formed by simpler molecular mechanisms than previously expected from the Ca2+ dynamics. Moreover, we analyzed how guanylyl cyclases and phosphodiesterases, which synthesize and degrade cGMP, respectively, contributed to cGMP and Ca2+ dynamics and thermotaxis behavior.

## Introduction

Perception of ambient temperature and subsequent modulation of behavior according to the ambient temperature are essential for animals to survive a fluctuating environment. *C. elegans* cultivated at certain temperature migrate toward that temperature on a thermal gradient (Hedgecock & Russell, 1975), which indicates that *C. elegans* can sense temperature, memorize past cultivation temperature, and compare the present temperature with the memorized past temperature. AFD, a major thermosensory neuron in *C. elegans* (Mori & Ohshima, 1995), increases intracellular Ca2+ level in response to temperature increase above around the cultivation temperature (Kimura *et al*, 2004). These properties of AFD are conserved even when AFD is disconnected from neural circuits in culture *in vitro* (Kobayashi *et al*, 2016), indicating that AFD can cell-autonomously sense temperature, memorize past cultivation temperature and compare the present temperature and the memorized past temperature.

In *C. elegans* AFD, three guanylyl cyclases (GCYs), GCY-8, GCY-18 and GCY-23, which are specifically expressed in AFD, and cyclic nucleotide-gated (CNG) channels TAX-4 and TAX-2, which are far more sensitive to cGMP than to cAMP, are essential for thermosensation (Inada *et al*, 2006; Komatsu *et al*, 1996; Ramot *et al*, 2008; Kobayashi *et al*, 2016; Komatsu *et al*, 1999): AFD in animals lacking all of three GCYs and those lacking TAX-4 or TAX-2 does not show any response to temperature stimuli. Since ectopic expression of these GCYs are sufficient to confer thermo-responsiveness to chemosensory neurons, which are otherwise irresponsive to temperature change, suggesting a possibility that these GCYs are temperature sensors (Takeishi *et al*, 2016). Although ion channels including transient receptor potential (TRP) channels are considered to play a major role in peripheral thermosensation in mammals (Vriens *et al*, 2014), it was recently shown that the subtype G of transmembrane guanylyl cyclase (GC-G) in the Grueneberg ganglion (GG) in the mouse nose is activated by cool temperature and necessary for cold-evoked behavior of mice (Chao *et al*, 2015). GC-G activates a cGMP-activated channel, CNGA3, which is proposed to mediate coolness-evoked response of GG neurons (Chao *et al*, 2015; Fleischer *et al*, 2009; Mamasuew *et al*, 2010). Thus, thermosensation by transmembrane guanylyl cyclases and subsequent activation of CNG channels might be a conserved mechanism of thermosensory transduction in different phyla.

Although cGMP apparently seems to mediate temperature information to Ca2+ increase in AFD neurons of *C. elegans*, whether and how cGMP level in AFD is temporally dynamic during temperature change has not been described. Therefore, we monitored cGMP level in AFD by expressing a genetically encoded cGMP probe, cGi500 (Russwurm *et al*, 2007). We found that cGMP level increased and decreased in response to temperature increase and decrease specifically at a sensory ending of AFD. Temperature at which cGMP started increasing and decreasing was dependent on past cultivation temperature similarly to Ca2+ dynamics. We also analyzed how GCYs and phophodiesterases (PDEs), which degrade cGMP, contribute to cGMP dynamics and Ca2+ dynamics in AFD and thermotaxis behavior.

## Results

### cGMP level in AFD dynamically responds to temperature change

To reveal whether and how cGMP level fluctuates in AFD, we expressed a genetically encoded fluorescent cGMP probe, cGi500, which increases CFP/YFP fluorescence ratio in response to increase of cGMP concentration (Russwurm *et al*, 2007), specifically in AFD and performed imaging analyses. CFP/YFP fluorescence ratio at the AFD sensory ending of wild type animals expressing cGi500 cultivated at 17°C or 23°C increased in response to temperature increase from around the past cultivation temperature and decreased in response to temperature decrease from around the past cultivation temperature, respectively (Figure 1A and B, left). This fluorescence ratio change was not observed in *gcy-23 gcy-8 gcy-18* triple mutant animals, in which AFD Ca2+ dynamics and thermotaxis behavior are abolished and therefore AFD cGMP dynamics are supposed to be abolished (Figure 1C). The seeming decrease and increase of CFP/YFP ratio in *gcy-23 gcy-8 gcy-18* triple mutant animals in response to temperature increase and decrease, respectively, are supposed to be derived from the effect of temperature change on the probe or the focus. Given the opposite trend in the increase and decrease in CFP/YFP ratio between wild type and *gcy-23 gcy-8 gcy-18* triple mutant animals, it is highly likely that the change of fluorescence ratio observed at the sensory endings of wild type animals reflects the genuine change of cGMP level.

**Figure 1.**
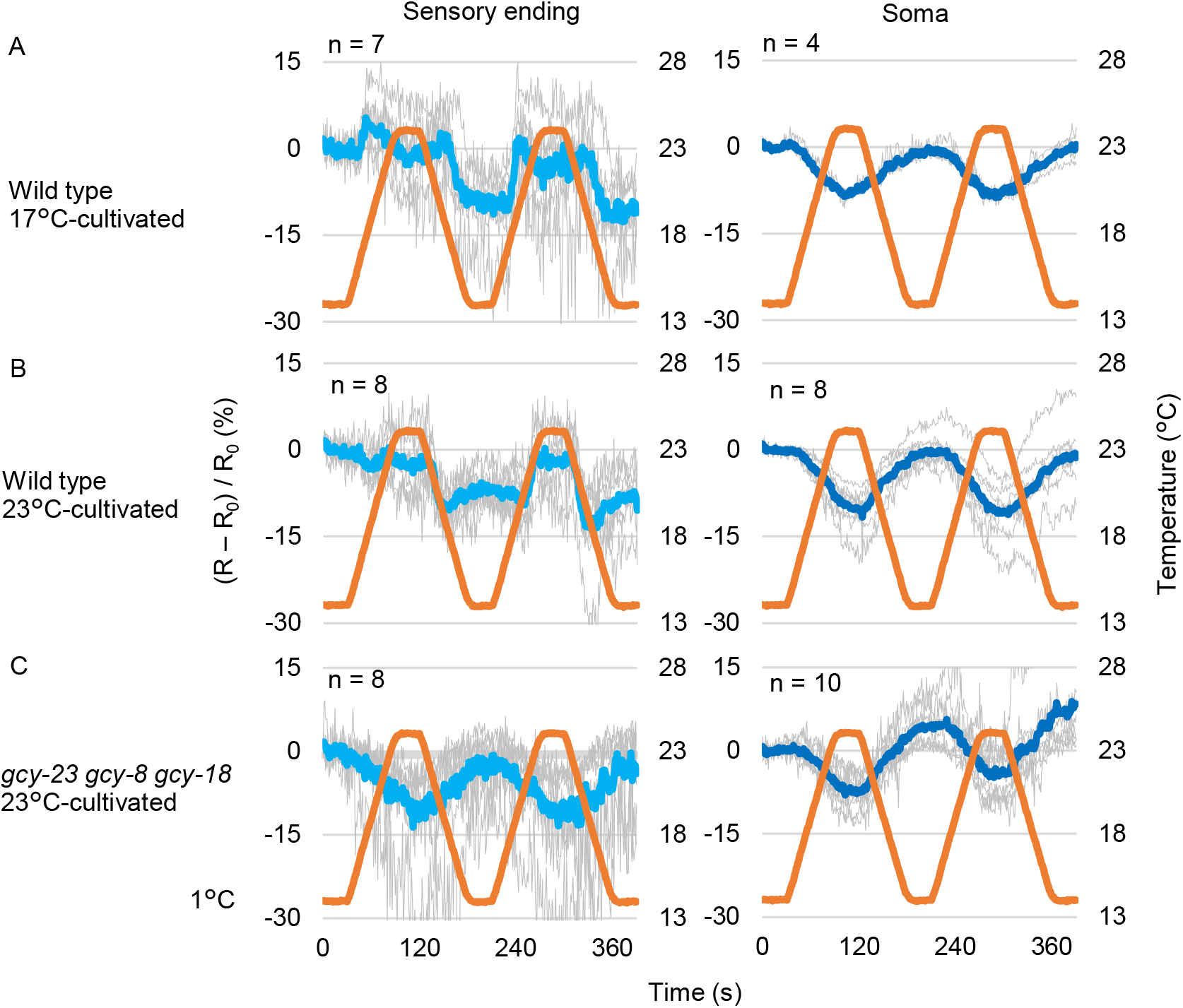
cGMP dynamics in AFD. A and B. Wild type animals expressing cGi500 cGMP indicator specifically in AFD thermosensory neurons (IK3110) were cultivated at 17°C (A) or 23°C (B). Blue and yellow fluorescence was monitored while temperature was increased from 14°C to 23°C and then decreased to 14°C, which were repeated twice as indicated (orange line). Temperature was increased and decreased by the rate of 1°C/6 sec. Individual (gray) and average fluorescence ratio (CFP/YFP) change at AFD sensory ending (blue) and soma (dark blue) is shown. C. *gcy-18 gcy-8 gcy-23* triple mutant animals expressing cGi500 cGMP indicator in AFD (IK3360) were cultivated at 23°C and subjected to imaging analysis. R_0_ is average of R (CFP/YFP) from t = 1 to t = 30.

In AFD soma, CFP/YFP fluorescence ratio seemed to decrease and increase in response to temperature increase and decrease, respectively, similarly to the AFD sensory ending in *gcy-23 gcy-8 gcy-18* triple mutants (Figure 1, right), suggesting that the cGMP level changes specifically at the sensory ending in AFD. These results are consistent with the sensory ending-specific localization of GCYs (Inada *et al*, 2006; Nguyen *et al*, 2014) and the fact that the laser-axotomized AFD sensory ending but not AFD soma changes Ca2+ level in response to temperature stimuli (Clark *et al*, 2006). Compartmentalized cGMP dynamics are also observed in olfactory sensory neuron AWC (Shidara *et al*, 2017) and O2-sensing neuron PQR (Couto *et al*, 2013).

Taken together, our results suggested that cGMP increases and decreases at the AFD sensory ending in response to temperature increase and decrease from around the past cultivation temperature, respectively. Although cGMP dynamics are supposed to be upstream of Ca2+ dynamics, the cGMP dynamics reflected not only the temperature change but also the comparison between the past and present temperature, suggesting that both the memory and the comparison are realized by simpler molecular mechanisms than previously expected from the Ca2+ dynamics.

### Guanylyl cyclases regulate onset temperature for change of cGMP level in AFD

Genetic analysis on thermotaxis behavior and AFD Ca2+ dynamics revealed that *gcy-8, gcy-18* and *gcy-23* are redundantly contribute to AFD thermo-receptivity (Inada *et al*, 2006; Takeishi *et al*, 2016). However, contribution of each *gcy* is not uniform since three of *gcy* double mutants, in which only one *gcy* gene out of the three is intact, exhibit differential abnormality in thermotaxis behavior (Inada *et al*, 2006; Wang *et al*, 2013)(Figure 2A) and AFD Ca2+ dynamics (Wasserman *et al*, 2011; Takeishi *et al*, 2016)(Supplementary Figure 1) especially when animals are cultivated at relatively high temperature such as 23°C. Namely, abnormality of *gcy-8 gcy-18* and *gcy23 gcy-8* mutants is the most and the least severe, respectively. We then examined how each GCY contributes to cGMP dynamics in AFD.

**Figure 2.**
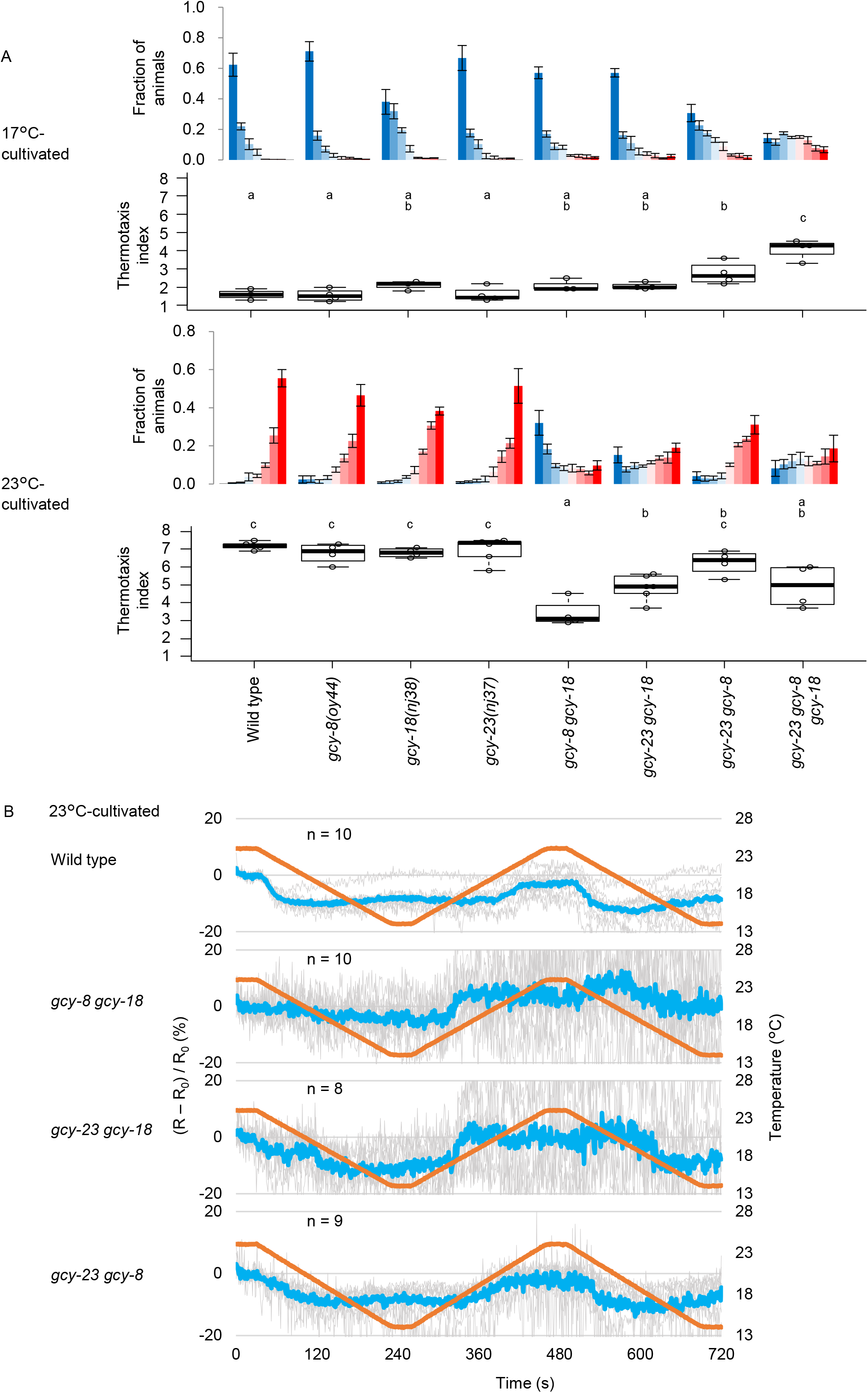
cGMP initiates increasing from lower temperature in *gcy* double mutants. A. Wild type animals and animals in which indicated *gcy* gene(s) is mutated were cultivated at 17°C or 23°C and then placed on a thermal gradient. The number of animals in each section of the thermal gradient was scored, and the proportion of animals in each section was plotted on histograms. n = 3 to 6 as indicated by open circles in boxplots. The error bars in histograms represent the standard error of mean (SEM). The thermotaxis indices were plotted on boxplots. The indices of strains marked with distinct alphabets differ significantly (p < 0.05) according to the Tukey-Kramer test. B. Wild type and indicated *gcy* double mutant animals that express cGi500 cGMP indicator in AFD were cultivated at 23°C and subjected to imaging analysis with temperature stimuli indicated (orange line). Temperature was increased and decreased by the rate of 1°C/20 sec. Individual (gray) and average (blue) fluorescence ratio (CFP/YFP) change at AFD sensory ending were plotted against time.

AFD in three *gcy* double mutant animals cultivated at 23°C exhibited lower onset temperature for cGMP increase and decrease than AFD in wild type animals (Figure 2B). The extent of abnormality was the most and the least severe in *gcy-8 gcy-18* and *gcy23 gcy-8* double mutants, respectively. These results were very well correlated to abnormality in the Ca2+ dynamics and the thermotaxis behavior; onset temperature for Ca2+ increase was the lowest and the highest (Figure S1), and the abnormality in thermotaxis was the most and the least severe (Figure 2A) in *gcy-8 gcy-18* and *gcy23 gcy-8* double mutants, respectively. Taken together, it seems that cGMP dynamics in AFD are composed of differential contribution from GCY-8, GCY-18 and GCY-23, and that Ca2+ level increases in response to cGMP increase. Interestingly, AFD Ca2+ level in animals cultivated at 23°C started decreasing already when temperature was constant (Supplementary Figure 1). This is in contrast with the temperature at which cGMP levels started decreasing, which was almost the same as the onset temperature for cGMP increase (Figure 2B). These results suggest that the AFD Ca2+ level is actively decreased regardless of cGMP level and are consistent with our previous finding that SLO K+ channels contribute to Ca2+ decrease in AFD after Ca2+ increase induced by temperature increase (Aoki *et al*, 2018); both SLO-1 and SLO-2 channels are activated by Ca2+ and the resulting K+ efflux may inactivate Ca2+ channels.

### Phosphodiesterases contribute to proper thermotaxis

cGMP is hydrolysed by phosphodiesterases (PDEs). Of six *C. elegans* PDEs, PDE-1, PDE-2, PDE-3 and PDE-5 are supposed to hydrolyse both cGMP and cAMP, whereas PDE-4 and PDE-6 are supposed to be cAMP-specific (Omori & Kotera, 2007; Lugnier, 2006; Liu *et al*, 2010). We aimed to analyze how these four PDEs affect cGMP and Ca2+ dynamics in AFD and thermotaxis behavior.

First, we analyzed thermotaxis of *pde* mutants. Of four mutant strains that lack single *pde* gene that hydrolyses cGMP, *pde-5* mutants were significantly defective in thermotaxis when cultivated at both 17°C and 23°C (Figure 3). *pde-1; pde-2* double mutants were also defective when cultivated at both 17°C and 23°C. Simultaneous mutation in *pde-5* and *pde-1* exhibited a synergistic effect especially when cultivated at 23°C. Effect of *pde-3* mutation on thermotaxis was marginal.

**Figure 3.**
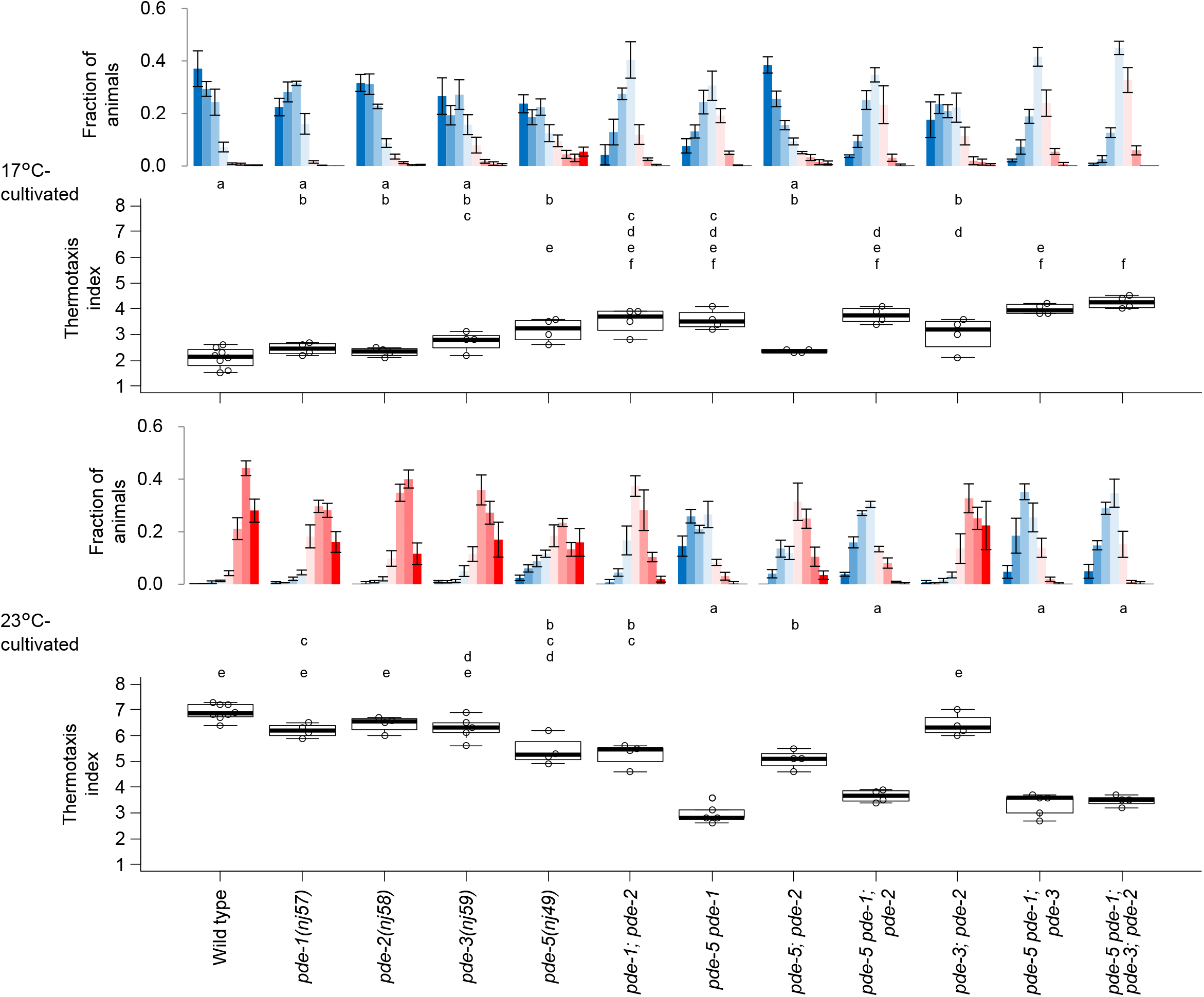
*pde* mutants are defective for thermotaxis behavior. Wild type and *pde* mutant animals indicated were cultivated at 17°C or 23°C and then subjected to thermotaxis assay. n = 4 to 8 as indicated by open circles in boxplots. The error bars in histograms represent the standard error of mean (SEM). The thermotaxis indices of strains marked with distinct alphabets differ significantly (p < 0.05) according to the Tukey-Kramer test.

### *pde-5* acts in AFD to regulate cGMP dynamics and thermotaxis behavior

*pde-5* is expressed in AFD (Wang *et al*, 2013). To examine whether *pde-5* acts in AFD to regulate cGMP dynamics and thermotaxis, we expressed *pde-5* specifically in AFD in *pde-5* mutants. AFD-specific *pde-5* expression rescued abnormality of thermotaxis of *pde-5* mutants cultivated at both 17°C and 23°C (Figure 4A), indicating that *pde-5* acts in AFD to regulate thermotaxis. We therefore monitored AFD cGMP dynamics in *pde-5* mutants. Although cGMP level is considered to increase in *pde-5* mutants, cGMP dynamics seemed abolished in *pde-5* mutants in animals cultivated at both 17°C and 23°C (Figure 4B). Abolished cGMP dynamics in *pde-5* mutants were rescued by AFD-specific expression of *pde-5* (Figure 4B), suggesting that *pde-5* cell-autonomously acts in AFD to regulate cGMP dynamics and thereby thermotaxis behavior. Expression of PDE-5 fused to GFP was observed throughout AFD including its sensory ending, indicating that PDE-5 may well affect cGMP dynamics at AFD sensory ending (Supplementary figure 2B).

**Figure 4.**
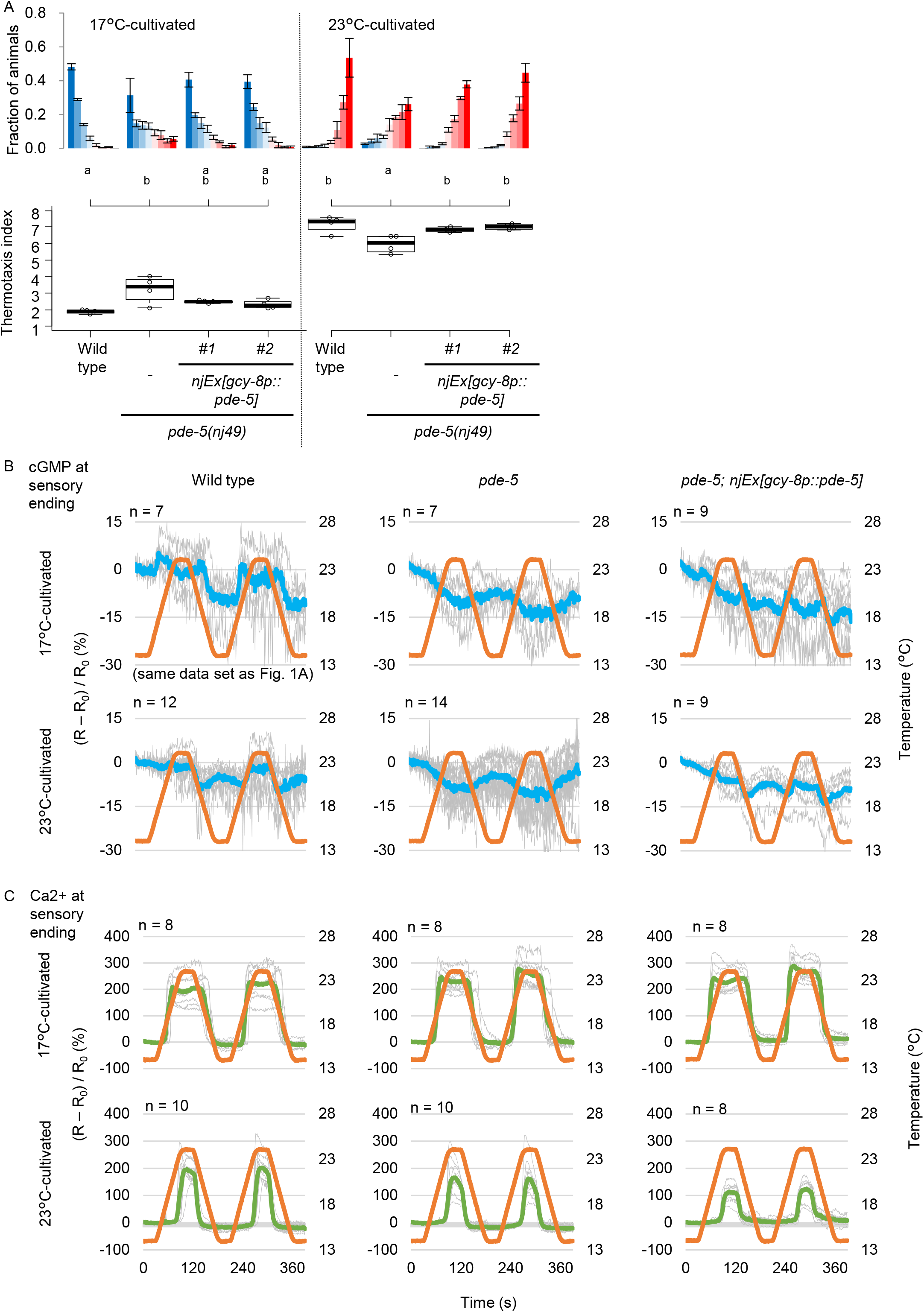
*pde-5* acts in AFD to regulate cGMP dynamics and thermotaxis. A. Wild type and *pde-5* animals and *pde-5* animals that express PDE-5 specifically in AFD were cultivated at 17°C or 23°C and then subjected to thermotaxis assay. n = 4. The error bars in histograms represent the standard error of mean (SEM). The thermotaxis indices of strains marked with distinct alphabets differ significantly (p < 0.05) according to the Tukey-Kramer test. B. Wild type and *pde-5* animals and *pde-5* mutant animals expressing PDE-5 in AFD that express cGi500 cGMP indicator in AFD were cultivated at 17°C or 23°C. Animals were then subjected to imaging analysis with temperature stimuli indicated (orange line). Temperature was increased and decreased by the rate of 1°C/6 sec. Individual (gray) and average (blue) fluorescence ratio (CFP/YFP) change at AFD sensory ending is shown. Data of wild type cultivated at 17°C are identical to those in Figure 1A. C. Wild type and *pde-5* animals and *pde-5* mutant animals expressing PDE-5 in AFD that express GCaMP3 Ca2+ indicator and tagRFP in AFD were cultivated at 17°C or 23°C. Animals were then subjected to imaging analysis with temperature stimuli indicated (orange line). Individual (gray) and average (pea green) fluorescence ratio (GCaMP/RFP) change at AFD sensory ending is shown.

We next aimed to examine the effect of abolished cGMP dynamics on Ca2+ dynamics in *pde-5* mutants. Surprisingly, the Ca2+ dynamics in *pde-5* mutants were indistinguishable from that in wild type (Figure 4C and Supplementary Figure 2). Given that the Ca2+ dynamics are supposed to be downstream of the cGMP dynamics, it seems paradoxical that the cGMP dynamics were abolished whereas the Ca2+ dynamics were intact in *pde-5* mutants. However, we have to note the possibilities that cGMP dynamics in *pde-5* mutants were out of cGi500’s operating range due to high basal cGMP level or small amplitude, and that the difference of Ca2+ dynamics that caused the abnormal thermotaxis was not prominent with the used temperature stimuli.

### *pde-1* and *pde-2* act in AFD to regulate thermotaxis

*pde-1* and *pde-2* are also expressed in AFD (Wang *et al*, 2013; Singhvi *et al*, 2016). AFD-specific expression of *pde-1* or *pde-2* partially rescued abnormality of *pde-1; pde-2* double mutants to the level of *pde-1* or *pde-2* single mutants (Figure 5A). These results indicate that both *pde-1* and *pde-2* act in AFD to regulate thermotaxis. We therefore monitored AFD cGMP and Ca2+ dynamics in *pde-1, pde-2* and *pde-1; pde-2* mutant animals. Surprisingly, both cGMP and Ca2+ dynamics in these mutants were indistinguishable from those in wild type (Figure 5B-D). Although it is possible that the difference of cGMP and Ca2+ dynamics that caused the abnormal thermotaxis were not prominent with the used temperature stimuli, *pde-1* and *pde-2* might possibly function downstream of Ca2+ increase to regulate thermotaxis.

**Figure 5.**
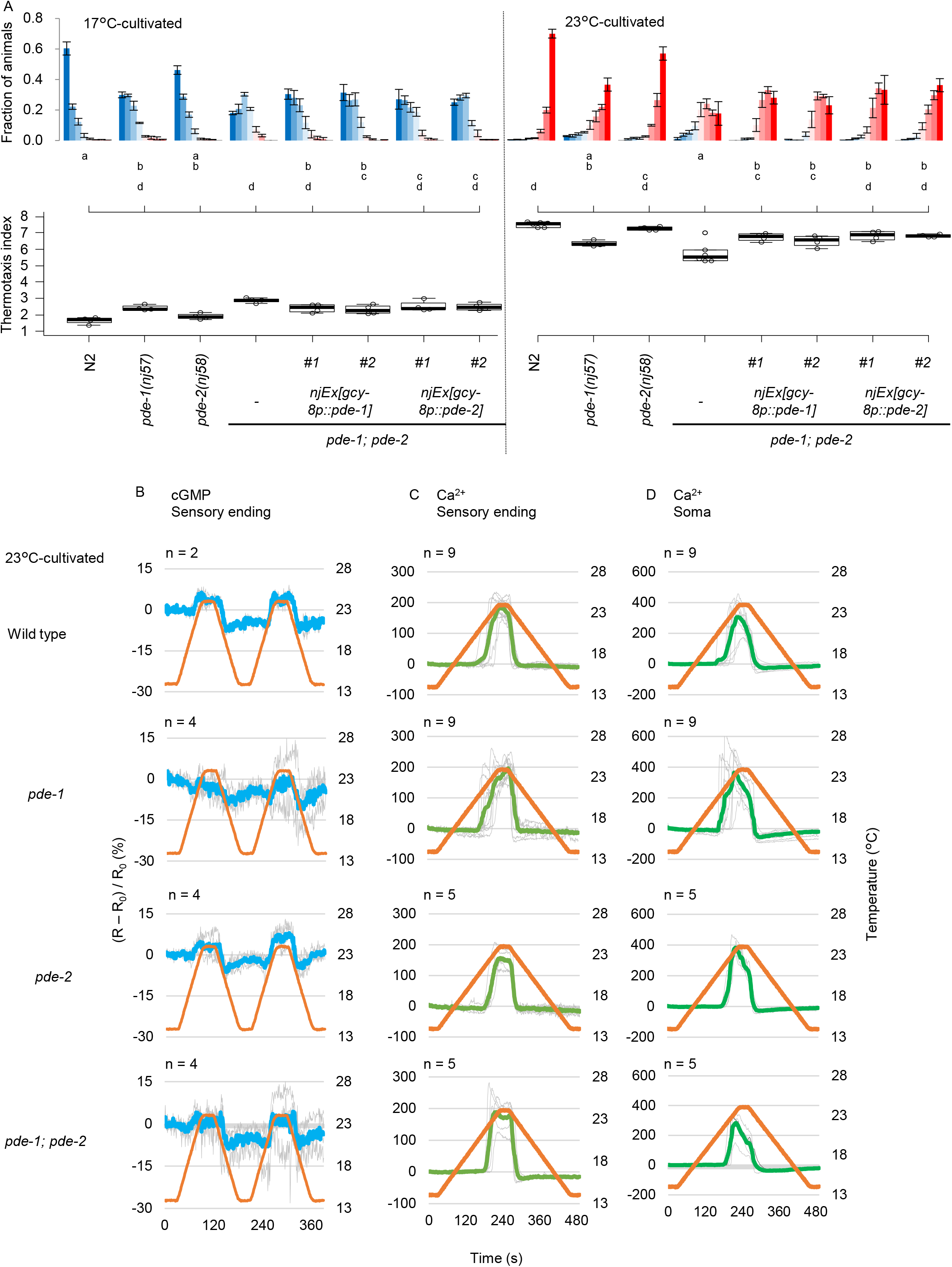
*pde-1*, *pde-2* act in AFD to regulate thermotaxis. A. Wild type, *pde-1, pde-2* and *pde-1; pde-2* animals and *pde-1; pde-2* animals that express PDE-1 or PDE-2 specifically in AFD were cultivated at 17°C or 23°C and then subjected to thermotaxis assay. n = 8 for N2 and *pde-1; pde-2*. n = 4 for others. The error bars in histograms represent the standard error of mean (SEM). The thermotaxis indices of strains marked with distinct alphabets differ significantly (p < 0.05) according to the Tukey-Kramer test. B. Wild type and mutant animals lacking *pde* gene(s) indicated that express cGi500 cGMP indicator in AFD were cultivated at 23°C and subjected to imaging analysis. Temperature was increased and decreased by the rate of 1°C/6 sec. Individual (gray) and average (blue) fluorescence ratio (CFP/YFP) change at AFD sensory ending is shown. C-D. Wild type and mutant animals lacking *pde* gene(s) indicated that express GCaMP3 Ca2+ indicator and tagRFP in AFD were cultivated at 23°C and subjected to imaging analysis. Temperature was increased and decreased by the rate of 1°C/20 sec. Individual (gray) and average (pea green or green) fluorescence ratio (GCaMP/RFP) change at AFD sensory ending (C) and soma (D) is shown.

### *pde-1* and *pde-5* synergize in AFD

We next examined whether the synergistic effect of *pde-1* and *pde-5* mutations on thermotaxis is derived from the act of these genes in AFD. AFD-specific expression of *pde-1* or *pde-5* partially rescued abnormality of *pde-5 pde-1* double mutants to the level of *pde-1* or *pde-5* single mutants (Figure 6A). cGMP dynamics in *pde-5 pde-1* double mutants were abolished similarly in *pde-5* mutants (Figure 6B). Interestingly, whereas Ca2+ dynamics in *pde-1* or *pde-5* single mutants were almost comparable to that in wild type, in *pde-5 pde-1* double mutants, Ca2+ gradually increased from lower temperature than the wild type and decreased gradually (Figure 6C, D). These results suggest that *pde-1* and *pde-5* redundantly function in AFD to suppress Ca2+ increase in response to temperature increase when ambient temperature is lower than cultivation temperature, to sharply decrease Ca2+ in response to temperature decrease, and therefore to limit the temperature range to which AFD responds, which is essential for proper thermotaxis.

**Figure 6.**
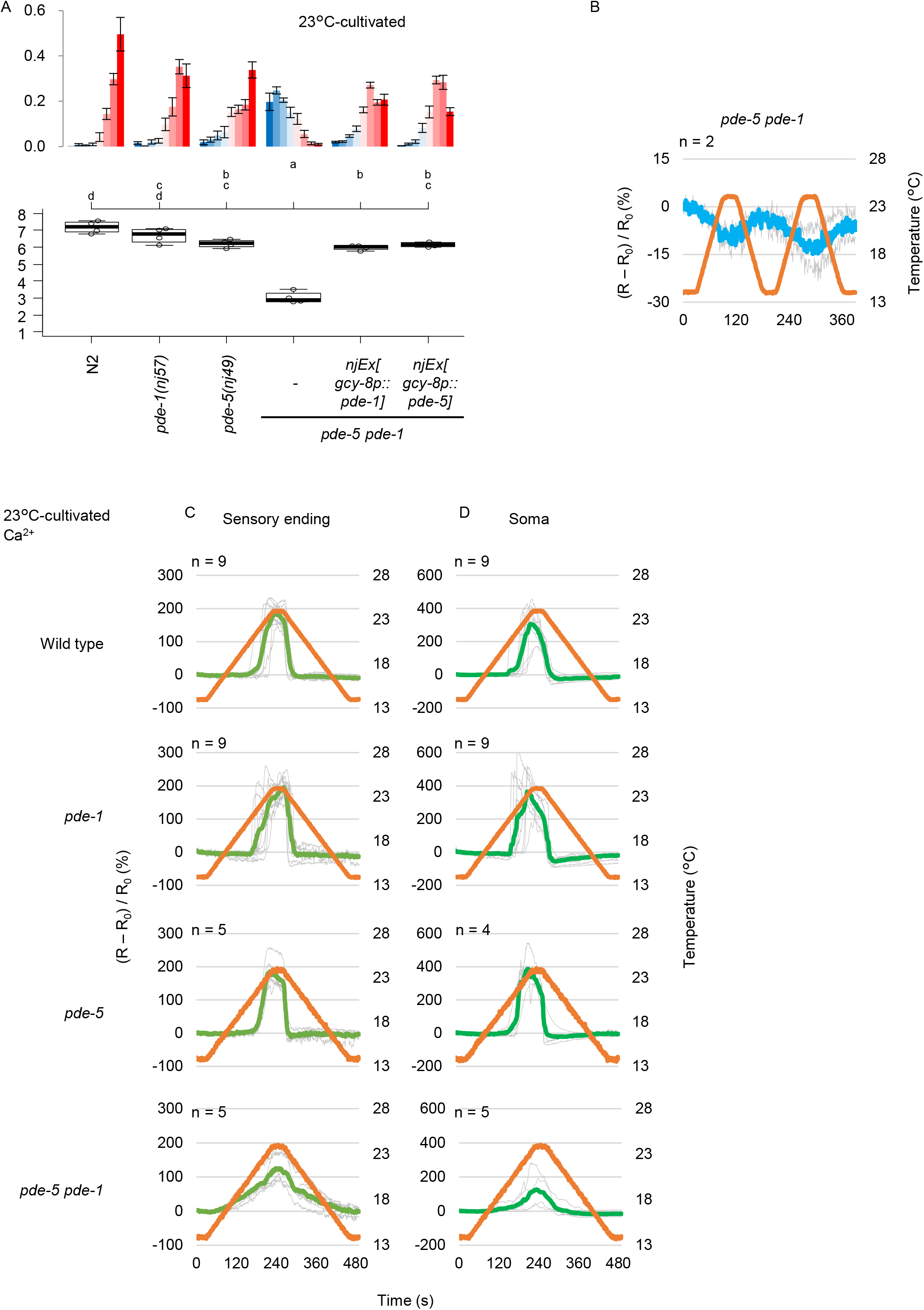
*pde-1* and *pde-5* synergize in AFD. A. Wild type, *pde-1*, *pde-5* and *pde-5 pde-1* double mutant animals and *pde-5 pde-1* animals that express *pde-1* or *pde-5* in AFD were cultivated at 23°C and then subjected to thermotaxis assay. n = 4. The error bars in histograms represent the standard error of mean (SEM). The thermotaxis indices of strains marked with distinct alphabets differ significantly (p < 0.05) according to the Tukey-Kramer test. B. *pde-5 pde-1* mutant animals that express cGi500 cGMP indicator in AFD were cultivated at 23°C and subjected to imaging analysis. C-D. Wild type, *pde-1*, *pde-5* and *pde-5 pde-1* double mutant animals that express GCaMP3 Ca2+ indicator and tagRFP in AFD were cultivated at 23°C and subjected to imaging analysis. Temperature was increased and decreased by the rate of 1°C/20 sec. Individual (gray) and average (pea green or green) fluorescence ratio (GCaMP/RFP) change at AFD sensory ending (C) and soma (D) is shown. Data of wild type and *pde-1* animals are identical to those in Figure 5C-D.

## Discussion

In this study, we demonstrated that cGMP increases and decreases in AFD thermosensory neurons of *C. elegans* in response to temperature increase and decrease, respectively. These cGMP dynamics were observed specifically at sensory endings but not at soma, which was in contrast to Ca2+ dynamics that are uniform among subcompartments in AFD such as sensory ending, dendrite, soma and axon. Given that GCYs and TAX-2 and TAX-4 CNG channels localize at the sensory ending in AFD and that Ca2+ dynamics are uniform all over AFD, there is supposed to be a mechanism that propagates Ca2+ increase from the sensory endings (Shindou *et al*, 2019). We also found that the cGMP dynamics, which are supposed to be upstream of Ca2+ dynamics, reflected the comparison between past cultivation temperature and present ambient temperature, indicating that AFD memorizes temperature by simpler molecular mechanisms than so far expected from the results of Ca2+ imaging and membrane potential recording. We further showed that AFD-specific GCYs determine onset temperature for cGMP increase, which is reflected on Ca2+ dynamics and thermotaxis behavior. Moreover, we described complex roles of PDEs in forming cGMP and Ca2+ dynamics and in thermotaxis behavior.

While cGMP and Ca2+ increase stereotypically in AFD in response to temperature increase from around the past cultivation temperature, how past cultivation temperature is memorized as onset temperature for cGMP and Ca2+ increase remains elusive. When GCY-23 and GCY-18 are ectopically expressed in chemosensory neurons, they confer low and high onset temperature regardless of animals’ cultivation temperature, respectively (Takeishi *et al*, 2016). However, in AFD of *gcy-23 gcy-8* double mutants, in which only GCY-18 is expressed out of the three GCYs essential for AFD thermosensation, onset temperature for Ca2+ increase changed according to cultivation temperature (Supplementary Figure 1). In line, the *gcy-23 gcy-8* double mutants change temperature preference in thermotaxis according to cultivation temperature in a similar manner as wild type animals (Figure 2). We showed in this study that cGMP dynamics, which are supposed to be upstream of Ca2+ dynamics, reflect past cultivation temperature. Taken together, these results raise two possibilities that can explain why AFD with only GCY-18 can adjust its onset temperature: First, a mechanism that assists GCY-18 to be activated at low temperature exists in AFD, or a mechanism that inhibits GCY-18 to be activated at low temperature in chemosensory neurons, the latter of which seems less likely. Second, given that expression level of *gcy*s change according to cultivation temperature (Yu *et al*, 2014; Aoki *et al*, 2018), regulation of GCY-18 expression level through *gcy-18* promoter, which was not involved during GCY-18 expression in chemosensory neurons (Takeishi *et al*, 2016), might be necessary for altering AFD onset temperature in *gcy-23 gcy-8* double mutants. Moreover, if three GCYs with different characteristics make heterodimers that have different characteristics from the homodimers, these variety of elements might contribute to fine-tune the responsiveness of AFD according to cultivation temperature.

The sensory ending of AFD has a unique and complicated morphology with microvilli, which might contribute to thermosensation (Singhvi *et al*, 2016; Bacaj *et al*, 2008), and the GCYs’ localization is restricted in the sensory ending including these microvilli by genes related to ciliary function (Nguyen *et al*, 2014; Inada *et al*, 2006). Loss of cilia during evolution is well correlated with loss of cGMP signaling pathway, suggesting that ciliary function and cGMP signaling are interdependent (Johnson & Leroux, 2010). Given that *C. elegans* mutants for the cilia-related genes are defective for thermotaxis (Nguyen *et al*, 2014; Tan *et al*, 2007), GCYs’ localization in the sensory ending may contribute to AFD’s ability to encode cultivation temperature with cGMP and Ca2+ dynamics and therefore to thermotaxis.

We showed that PDE-1, PDE-2 and PDE-5 act in AFD to regulate thermotaxis. However, contributions of PDEs to cGMP and Ca2+ dynamics and to thermotaxis did not seem so straight-forward as those of GCYs. How these PDEs, especially PDE-5, affect cGMP dynamics needs to be re-evaluated once a more sensitive cGMP indicator is developed. However, we at least found PDE-1 and PDE-5 redundantly function to restrict the dynamic range of AFD’s Ca2+ response, which seems to be essential for proper thermotaxis behavior.

## Materials and Methods

### Experimental model and subject details

*C. elegans* strains were cultivated on nematode growth medium (NGM) plates seeded with *E. Coli* OP50 strain (Caenorhabditis Genetics Center (CGC), Twin Cities, MN, USA) as described (Brenner, 1974). N2 (Bristol) was used as the wild type strain unless otherwise indicated. Transgenic lines were generated by injecting plasmid DNA directly into hermaphrodite gonad as described (Mello *et al*, 1991). Strains used in this study were listed in Supplementary Table 1.

### Behavioral assays

Population thermotaxis (TTX) assays were performed as described previously (Ito *et al*, 2006). Briefly, 50 to 250 animals cultivated at 17°C or 23°C were placed on the center of the assay plates without food with the temperature gradient of 17–23°C and were allowed to freely move for 1 h. The assay plate was divided into eight sections along the temperature gradient, and the number of the adult animals in each section was scored. Ratio of animal numbers in each section was plotted in histograms. Thermotaxis indices were calculated as shown below:

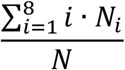

N_i_: number of animals in each section i (i = 1 to 8), N: total number of animals on the test plate.

### Plasmids

pAF-EXPR-26 *gcy-37p::cGi500::unc-54 3’UTR* was a gift from Dr. de Bono (Couto *et al*, 2013). cGi500 was sub-cloned into pDONR221 using the Gateway BP reaction (Thermo). To create a plasmid to express cGi500 in AFD, gcy-8 promoter, cGi500 cDNA and *unc-54* 3’UTR were fused by MultiSite Gateway Technology (Thermo Fisher Scientific, Waltham, MA, USA).

*pde-1b, pde-2a* and *pde-5* cDNAs were PCR amplified from DupLEX-A Yeast Two-Hybrid cDNA library *C. elegans* (adult) (OriGene, Rockville, MD) and cloned into AgeI-NotI sete of pIA138 *gcy-8p::VN173::unc-54 3’UTR*.

Plasmids used in this study are listed in Supplementary Table 2. Details regarding the plasmid constructs including sequences can be obtained from the authors.

### Imaging analyses

cGMP and Calcium imaging was performed as described elsewhere (Kobayashi *et al*, 2016; Aoki *et al*, 2018). Briefly, a single adult animal that expressed genetically encoded cGMP indicator cGi500(ref) or that express calcium indicator GCaMP3 (Tian *et al*, 2009) and tagRFP in AFD was placed on a 10% agar pad on a cover slip with 0.1 μm polystyrene beads (Polysciences, Warrington, PA, USA) and covered by another cover slip for immobilization (Kim *et al*, 2013). The immobilized animals were placed on a Peltier-based temperature controller (Tokai Hit, Fujinomiya, Japan) on a stage of BX61WI microscope (Olympus, Tokyo, Japan). The cyan and yellow, or red and green fluorescence was separated by the Dual-View optics system (Molecular Devices, Sunnyvale, CA, USA), and the images were captured by an EM-CCD camera C9100-13 ImageEM (Hamamatsu Photonics, Japan) at 1 Hz frame rate. Excitation pulses were generated by SPECTRA light engine (Lumencor, Beaverton, OR, USA). The fluorescence intensities were measured by the MetaMorph imaging system (Molecular Devices). Change of fluorescence ratio R (CFP/YFP for cGMP imaging by cGi500 and GFP/RFP for Ca2+ imaging by GCaMP3 and tagRFP), (R – R_0_) / R_0_, was plotted, where R_0_ is average of R from t = 1 to t = 30.

Expression of PDE-5::GFP was observed with LSM880 confocal microscope (Zeiss, Oberkochen, Germany).

### Statistical Analysis

The error bars in histograms for thermotaxis assays indicate the standard error of mean (SEM). In the boxplots, the bottom and top of boxes represent the first and third quartiles, and the band inside the box represents the median. The ends of the lower and upper whiskers represent the lowest datum still within the 1.5 interquartile range (IQR), which is equal to the difference between the third and first quartiles, of the lower quartile, and the highest datum still within the 1.5 IQR of the upper quartile, respectively. For multiple-comparison tests, one-way analyses of variance (ANOVAs) were performed, followed by Tukey-Kramer test. Static analyses were done by R.

## Data Availability

All strains and plasmids generated in this study are available upon request. One supplemental figure, a plasmid list and a strain list are provided.

## Acknowledgement

We thank H. Matsuyama for critical reading. Some strains were provided by the *Caenorhabditis* Genetics Center (CGC), which is funded by NIH Office of Research Infrastructure Programs (P40 OD010440). This work was supported by JSPS KAKENHI Grant Numbers JP 16H01272, JP 16H02516 and JP 18H04693.

## Author contributions

I.A. designed and performed experiments, wrote the manuscript, and supervised.

M.S. performed experiments.

S. N. designed experiments.

I.M. supervised and secured funding.

---

**Figure S1.**
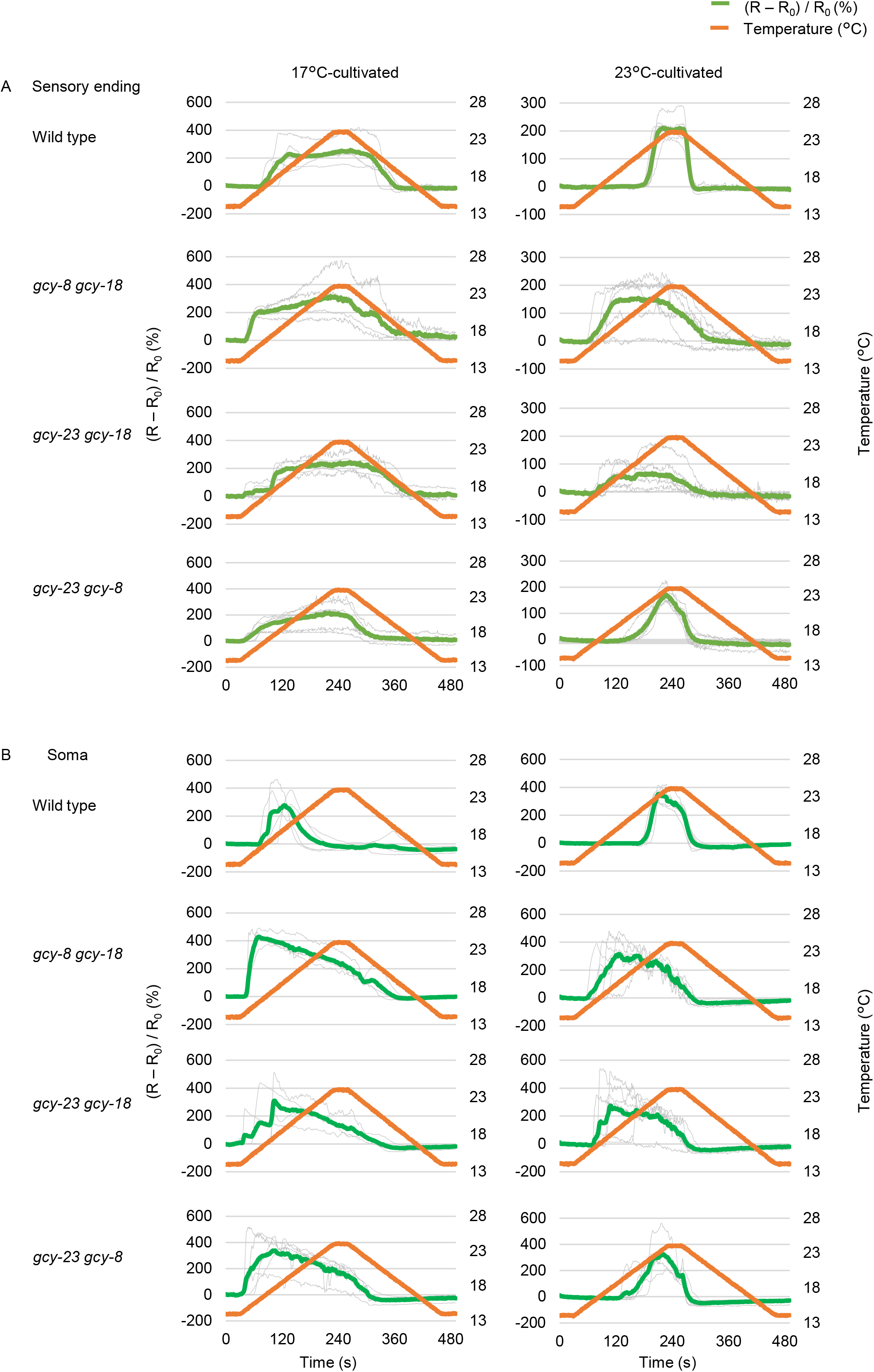
Ca2+ initiates increasing from lower temperature in *gcy* double mutants (related to Figure 2) Wild type and *gcy* double mutant animals indicated that express GCaMP3 Ca2+ indicator and tagRFP in AFD were cultivated at 17°C (left) or 23°C (right) and subjected to imaging analysis with temperature stimuli indicated (orange line). Temperature was increased and decreased by the rate of 1°C/20 sec. n = 4 to 6. Individual (gray) and average fluorescence ratio (GCaMP/RFP) change at AFD sensory ending (A, pea green) and soma (B, green) is shown.

**Figure S2.**
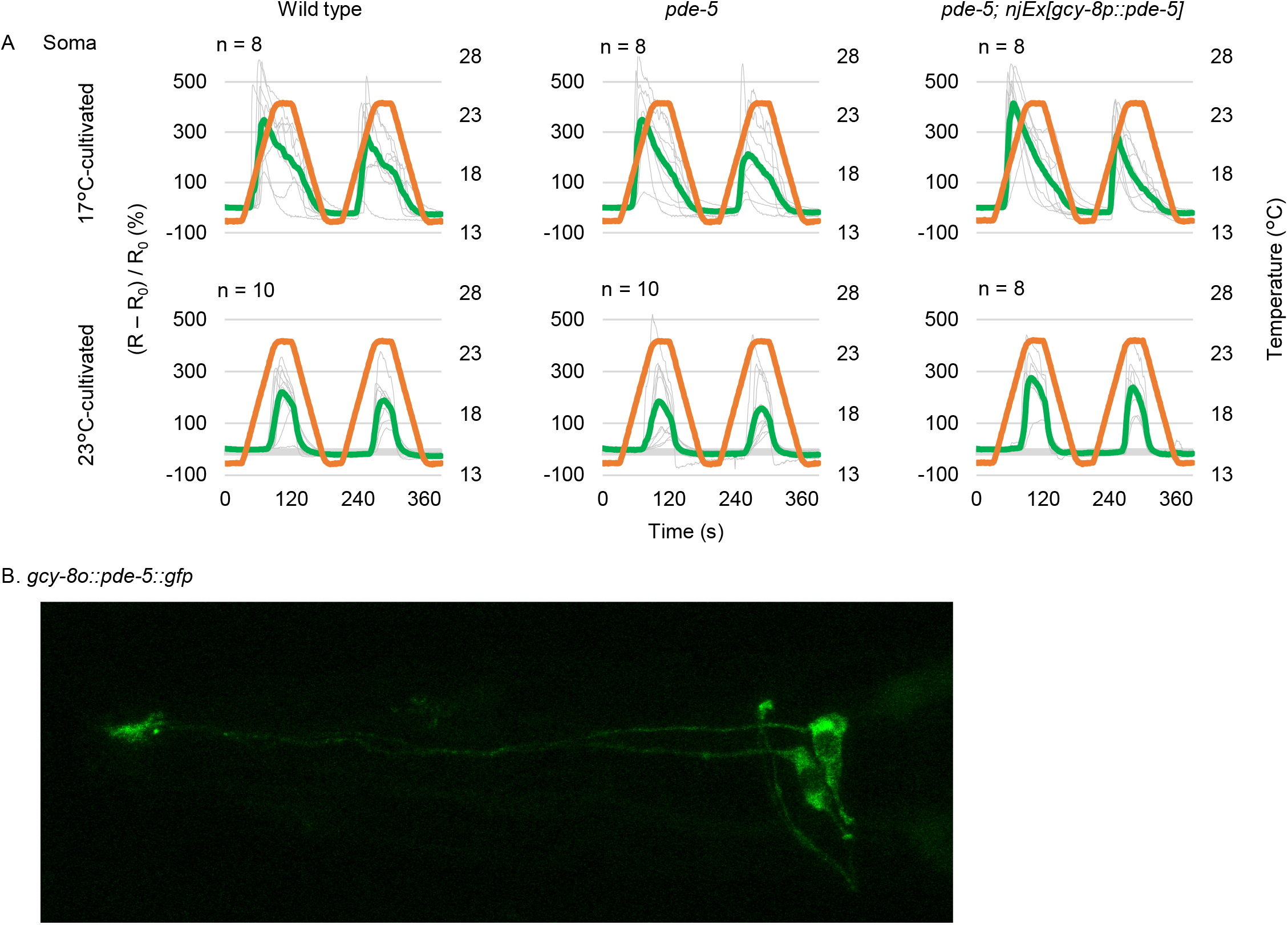
Ca2+ imaging of *pde-5* mutants at soma (related to Figure 4) A. Wild type and *pde-5* animals and *pde-5* mutant animals expressing PDE-5 in AFD that express GCaMP3 Ca2+ indicator and tagRFP in AFD were cultivated at 17°C or 23°C. Animals were then subjected to imaging analysis with temperature stimuli indicated (orange line). Individual (gray) and average (green) fluorescence ratio (GCaMP/RFP) change at AFD soma is shown. B. *pde-5(nj49); njEx1414[gcy-8p::pde-5::GFP, ges-1p::NLStagRFP]* was subjected to microscopic analysis with Zeiss LSM88 confocal microscopy.

**Supplementary Table 1.** Strain list.

| Description | Source | Identifier |
| --- | --- | --- |
| *C. elegans*: wild isolate | CGC | WormBase: N2<br>RRID:WB-STRAIN:N2_(ancestral) |
| *pde-1*(nj57) |  | IK0616 |
| *pde-2*(nj58) |  | IK0618 |
| *pde-3*(nj59) |  | IK0620 |
| *pde-5*(nj49) |  | IK0510 |
| *pde-1*(nj57); *pde-2*(nj58) |  | IK0792 |
| *pde-5*(nj49) *pde-1*(nj57) |  | IK0794 |
| *pde-5*(nj49); *pde-2*(nj58) |  | IK0796 |
| *pde-5*(nj49) *pde-1*(nj57); *pde-2*(nj58) |  | IK0798 |
| *pde-3*(nj59); *pde-2*(nj58) |  | IK1395 |
| *pde-5*(nj49) *pde-1*(nj57); *pde-3*(nj59) |  | IK1400 |
| *pde-5*(nj49) *pde-1*(nj57); *pde-3*(nj59); *pde-2*(nj58) |  | IK1401 |
| *pde-5*(nj49); njEx1408[gcy-8p::pde-5 cDNA, ges-1p::GFP] line1 |  | IK3352 |
| *pde-5*(nj49); njEx1409[gcy-8p::pde-5 cDNA, ges-1p::GFP] line2 |  | IK3353 |
| njEx1285[gcy-8p::cGi500(60 ng/ul), ges-1p::NLStagRFP] line1 |  | IK3110 |
| *pde-5*(nj49); njEx1285[gcy-8p::cGi500, ges-1p::NLStagRFP] line1 |  | IK3410 |
| *pde-5*(nj49); njEx1285[gcy-8p::cGi500, ges-1p::NLStagRFP]; njEx1408[gcy-8p::pde-5 cDNA, ges-1p::GFP] |  | IK3439 |
| njEx359[gcy-8p::TagRFP 20ng, gcy-8p::GCaMP3 50ng]] |  | IK0855 |
| *pde-5*(nj49); njEx359[gcy-8p::TagRFP 20ng, gcy-8p::GCaMP3 50ng]] |  | IK3434 |
| *pde-5*(nj49); njEx359[gcy-8p::TagRFP 20ng, gcy-8p::GCaMP3 50ng]]; njEx1408[gcy-8p::pde-5 cDNA, ges-1p::GFP] |  | IK3451 |
| *pde-5*(nj49); njEx1414[gcy-8p::pde-5::GFP, ges-1p::NLStagRFP] line1 |  | IK3358 |
| *pde-1*(nj57); *pde-2*(nj58); njEx1410[gcy-8p::pde-1 cDNA, ges-1p::GFP] line1 |  | IK3354 |
| *pde-1*(nj57); *pde-2*(nj58); njEx1411[gcy-8p::pde-1 cDNA, ges-1p::GFP] line2 |  | IK3355 |
| *pde-1*(nj57); *pde-2*(nj58); njEx1412[gcy-8p::pde-2 cDNA, ges-1p::GFP] line1 |  | IK3356 |
| *pde-1*(nj57); *pde-2*(nj58); njEx1413[gcy-8p::pde-2 cDNA, ges-1p::GFP] line2 |  | IK3357 |
| *pde-1*(nj57); njEx1285[gcy-8p::cGi500, ges-1p::NLStagRFP] line1 |  | IK3407 |
| *pde-2*(nj58); njEx1285[gcy-8p::cGi500, ges-1p::NLStagRFP] |  | IK3409 |
| *pde-1*(nj57); *pde-2*(nj58); njEx1285[gcy-8p::cGi500, ges-1p::NLStagRFP] |  | IK3433 |
| *pde-5*(nj49) *pde-1*(nj57); njEx1285[gcy-8p::cGi500, ges-1p::NLStagRFP] |  | IK3440 |
| *pde-5*(nj49) *pde-1*(nj57); njEx359[gcy-8p::TagRFP 20ng, gcy-8p::GCaMP3 50ng]] |  | IK3453 |
| *pde-1*(nj57); njEx359[gcy-8p::TagRFP 20ng, gcy-8p::GCaMP3 50ng]] |  | IK3452 |
| *pde-5*(nj49) *pde-1*(nj57); njEx1410[gcy-8p::pde-1 cDNA, ges-1p::GFP] |  | IK3457 |
| *pde-5*(nj49) *pde-1*(nj57); njEx1408[gcy-8p::pde-5 cDNA, ges-1p::GFP] |  | IK3458 |
| gcy-23(nj37) gcy-8(oy44) gcy-18(nj38); njEx1285[gcy-8p::cGi500, ges-1p::NLStagRFP] |  | IK3360 |
| gcy-8(oy44) gcy-18(nj38); njEx1285[gcy-8p::cGi500, ges-1p::NLStagRFP] |  | IK3365 |
| gcy-23(nj37) gcy-8(oy44); njEx1285[gcy-8p::cGi500, ges-1p::NLStagRFP] |  | IK3366 |
| gcy-23(nj37) gcy-18(nj38); njEx1285[gcy-8p::cGi500, ges-1p::NLStagRFP] |  | IK3367 |
| gcy-8(oy44) gcy-18(nj38); njEx359[gcy-8p::TagRFP 20ng, gcy-8p::GCaMP3 50ng]] |  | IK3454 |
| gcy-23(nj37) gcy-8(oy44); njEx359[gcy-8p::TagRFP 20ng, gcy-8p::GCaMP3 50ng]] |  | IK3455 |
| gcy-23(nj37) gcy-18(nj38); njEx359[gcy-8p::TagRFP 20ng, gcy-8p::GCaMP3 50ng]] |  | IK3456 |
| tax-4(p678); njEx1285[gcy-8p::cGi500, ges-1p::NLStagRFP] |  | IK3412 |
| *pde-1*(nj57); *pde-2*(nj58); njEx1404[snb-1p::pde-1 cDNA, ges-1p::GFP] line1 |  | IK3348 |
| *pde-1*(nj57); *pde-2*(nj58); njEx1405[snb-1p::pde-1 cDNA, ges-1p::GFP] line2 |  | IK3349 |
| *pde-1*(nj57); *pde-2*(nj58); njEx1406[snb-1p::pde-2 cDNA, ges-1p::GFP] line1 |  | IK3350 |
| *pde-1*(nj57); *pde-2*(nj58); njEx1407[snb-1p::pde-2 cDNA, ges-1p::GFP] line2 |  | IK3351 |
| gcy-8(oy44); njEx1285[gcy-8p::cGi500, ges-1p::NLStagRFP] |  | IK3361 |
| gcy-18(nj38); njEx1285[gcy-8p::cGi500, ges-1p::NLStagRFP] |  | IK3362 |
| gcy-23(nj37); njEx1285[gcy-8p::cGi500, ges-1p::NLStagRFP] line1 |  | IK3363 |
| gcy-23(nj37); njEx1285[gcy-8p::cGi500, ges-1p::NLStagRFP] line2 |  | IK3364 |

**Supplementary Table 2.** Plasmid list.

| Description | Source | Identifier |
| --- | --- | --- |
| *gcy-8p::cGi500* | This paper | pMS001 |
| *gcy-8p::pde-1b* | This paper | pIA140 |
| *gcy-8p::pde-2a* | This paper | pIA141 |
| *gcy-8p::pde-5* | This paper | pIA143 |
| *gcy-8p::pde-5::gfp* | This paper | pMS005 |

## References

Aoki I, Tateyama M, Shimomura T, Ihara K, Kubo Y, Nakano S & Mori I (2018) SLO potassium channels antagonize premature decision making in C. elegans. Commun. Biol. 1: 123 Available at: http://www.nature.com/articles/s42003-018-0124-5

Bacaj T, Tevlin M, Lu Y & Shaham S (2008) Glia Are Essential for Sensory Organ Function in C. elegans. Science (80-.). 322: 744–747 Available at: http://www.sciencemag.org/cgi/doi/10.1126/science.1163074

Brenner S (1974) The genetics of Caenorhabditis elegans. Genetics 77: 71–94

Chao Y-C, Chen C-C, Lin Y-C, Breer H, Fleischer J & Yang R-B (2015) Receptor guanylyl cyclase-G is a novel thermosensory protein activated by cool temperatures. EMBO J. 34: 294–306 Available at: http://www.pubmedcentral.nih.gov/articlerender.fcgi?artid=4339118&tool=pmcentrez&rendertype=abstract

Clark DA, Biron D, Sengupta P & Samuel ADT (2006) The AFD Sensory Neurons Encode Multiple Functions Underlying Thermotactic Behavior in Caenorhabditis elegans. J. Neurosci. 26: 7444–7451 Available at: http://www.jneurosci.org/cgi/doi/10.1523/JNEUROSCI.1137-06.2006

Couto A, Oda S, Nikolaev VO, Soltesz Z & de Bono M (2013) In vivo genetic dissection of O2-evoked cGMP dynamics in a Caenorhabditis elegans gas sensor. Proc. Natl. Acad. Sci. U. S. A. 110: E3301–10 Available at: http://www.pubmedcentral.nih.gov/articlerender.fcgi?artid=3761592&tool=pmcentrez&rendertype=abstract

Fleischer J, Mamasuew K & Breer H (2009) Expression of cGMP signaling elements in the Grueneberg ganglion. Histochem. Cell Biol. 131: 75–88 Available at: http://link.springer.com/10.1007/s00418-008-0514-8 [Accessed June 25, 2019]

Hedgecock EM & Russell RL (1975) Normal and mutant thermotaxis in the nematode Caenorhabditis elegans. Proc. Natl. Acad. Sci. U. S. A. 72: 4061–5 Available at: http://www.pubmedcentral.nih.gov/articlerender.fcgi?artid=433138&tool=pmcentrez&rendertype=abstract

Inada H, Ito H, Satterlee J, Sengupta P, Matsumoto K & Mori I (2006) Identification of guanylyl cyclases that function in thermosensory neurons of Caenorhabditis elegans. Genetics 172: 2239–52 Available at: http://www.ncbi.nlm.nih.gov/pubmed/16415369 [Accessed February 7, 2013]

Ito H, Inada H & Mori I (2006) Quantitative analysis of thermotaxis in the nematode Caenorhabditis elegans. J. Neurosci. Methods 154: 45–52

Johnson J-LF & Leroux MR (2010) cAMP and cGMP signaling: sensory systems with prokaryotic roots adopted by eukaryotic cilia. Trends Cell Biol. 20: 435–444 Available at: https://www.sciencedirect.com/science/article/pii/S096289241000098X?via%3Dihub [Accessed November 16, 2018]

Kim E, Sun L, Gabel C V. & Fang-Yen C (2013) Long-Term Imaging of Caenorhabditis elegans Using Nanoparticle-Mediated Immobilization. PLoS One 8:

Kimura KD, Miyawaki A, Matsumoto K & Mori I (2004) The C. elegans thermosensory neuron AFD responds to warming. Curr. Biol. 14: 1291–1295 Available at: http://dx.doi.org/10.1016/j.cub.2004.06.060 [Accessed May 16, 2013]

Kobayashi K, Nakano S, Amano M, Tsuboi D, Nishioka T, Ikeda S, Yokoyama G, Kaibuchi K & Mori I (2016) Single-Cell Memory Regulates a Neural Circuit for Sensory Behavior. Cell Rep. 14: 11–21 Available at: http://dx.doi.org/10.1016/j.celrep.2015.11.064

Komatsu H, Jin YH, L’Etoile ND, Mori I, Bargmann CI, Akaike N & Ohshima Y (1999) Functional reconstitution of a heteromeric cyclic nucleotide-gated channel of Caenorhabditis elegans in cultured cells. Brain Res. 821: 160–168 Available at: http://www.ncbi.nlm.nih.gov/pubmed/10064800

Komatsu H, Mori I, Rhee JS, Akaike N & Ohshima Y (1996) Mutations in a cyclic nucleotide-gated channel lead to abnormal thermosensation and chemosensation in C. elegans. Neuron 17: 707–718 Available at: http://www.ncbi.nlm.nih.gov/pubmed/8893027

Liu J, Ward A, Gao J, Dong Y, Nishio N, Inada H, Kang L, Yu Y, Ma D, Xu T, Mori I, Xie Z & Xu XZS (2010) C. elegans phototransduction requires a G protein–dependent cGMP pathway and a taste receptor homolog. Nat. Neurosci. 13: 715–722 Available at: http://www.nature.com/articles/nn.2540 [Accessed December 2, 2018]

Lugnier C (2006) Cyclic nucleotide phosphodiesterase (PDE) superfamily: A new target for the development of specific therapeutic agents. Pharmacol. Ther. 109: 366–398 Available at: https://www.sciencedirect.com/science/article/pii/S0163725805001580?via%3Dihub [Accessed November 24, 2018]

Mamasuew K, Michalakis S, Breer H, Biel M & Fleischer J (2010) The cyclic nucleotide-gated ion channel CNGA3 contributes to coolness-induced responses of Grueneberg ganglion neurons. Cell. Mol. Life Sci. 67: 1859–1869 Available at: http://link.springer.com/10.1007/s00018-010-0296-8 [Accessed June 25, 2019]

Mello CC, Kramer JM, Stinchcomb D & Ambros V (1991) Efficient gene transfer in C. elegans: extrachomosomal maintenance and integration of transforming sequences. EMBO J. 10: 3959–3970 Available at: http://www.pubmedcentral.nih.gov/articlerender.fcgi?artid=453137&tool=pmcentrez&rendertype=abstract

Mori I & Ohshima Y (1995) Neural regulation of thermotaxis in Caenorhabditis elegans. Nature 376: 344–348 Available at: http://dx.doi.org/10.1038/376344a0

Nguyen PAT, Liou W, Hall DH & Leroux MR (2014) Ciliopathy proteins establish a bipartite signaling compartment in a C. elegans thermosensory neuron. J. Cell Sci. 127: 5317–30 Available at: http://jcs.biologists.org/cgi/doi/10.1242/jcs.157610

Omori K & Kotera J (2007) Overview of PDEs and Their Regulation. Circ. Res. 100: 309–327 Available at: https://www.ahajournals.org/doi/10.1161/01.RES.0000256354.95791.f1 [Accessed November 21, 2018]

Ramot D, MacInnis BL & Goodman MB (2008) Bidirectional temperature-sensing by a single thermosensory neuron in C. elegans. Nat. Neurosci. 11: 908–15 Available at: http://dx.doi.org/10.1038/nn.2157 [Accessed February 6, 2014]

Russwurm M, Mullershausen F, Friebe A, Jäger R, Russwurm C & Koesling D (2007) Design of fluorescence resonance energy transfer (FRET)-based cGMP indicators: a systematic approach. Biochem. J. 407: 69–77 Available at: http://biochemj.org/lookup/doi/10.1042/BJ20070348

Shidara H, Hotta K & Oka K (2017) Compartmentalized cGMP Responses of Olfactory Sensory Neurons in Caenorhabditis elegans. J. Neurosci. 37: 3753–3763 Available at: http://www.ncbi.nlm.nih.gov/pubmed/19889851%5Cnhttp://jn.physiology.org/content/103/1/441.full.pdf [Accessed May 13, 2014]

Shindou T, Ochi-Shindou M, Murayama T, Saita E, Momohara Y, Wickens JR & Maruyama IN (2019) Active propagation of dendritic electrical signals in C. elegans. Sci. Rep. 9: 3430 Available at: http://www.nature.com/articles/s41598-019-40158-9 [Accessed September 2, 2019]

Singhvi A, Liu B, Friedman CJ, Fong J, Lu Y, Huang XY, Shaham S, Singhvi A, Liu B, Friedman CJ, Fong J, Lu Y & Huang XY (2016) A Glial K/Cl Transporter Controls Neuronal Receptive Ending Shape by Chloride Inhibition of an rGC. Cell 165: 936–948 Available at: http://dx.doi.org/10.1016/j.cell.2016.03.026

Takeishi A, Yu Y V., Hapiak VM, Bell HW, O’Leary T & Sengupta P (2016) Receptor-type Guanylyl Cyclases Confer Thermosensory Responses in C. elegans. Neuron 90: 235–244 Available at: http://dx.doi.org/10.1016/j.neuron.2016.03.002

Tan PL, Barr T, Inglis PN, Mitsuma N, Huang SM, Garcia-Gonzalez MA, Bradley BA, Coforio S, Albrecht PJ, Watnick T, Germino GG, Beales PL, Caterina MJ, Leroux MR, Rice FL & Katsanis N (2007) Loss of Bardet Biedl syndrome proteins causes defects in peripheral sensory innervation and function. Proc. Natl. Acad. Sci. 104: 17524–17529 Available at: http://www.ncbi.nlm.nih.gov/pubmed/10468634 [Accessed November 16, 2018]

Tian L, Hires SA, Mao T, Huber D, Chiappe ME, Chalasani SH, Petreanu L, Akerboom J, McKinney SA, Schreiter ER, Bargmann CI, Jayaraman V, Svoboda K & Looger LL (2009) Imaging neural activity in worms, flies and mice with improved GCaMP calcium indicators. Nat. Methods 6: 875–881 Available at: http://www.nature.com/doifinder/10.1038/nmeth.1398 [Accessed September 25, 2013]

Vriens J, Nilius B & Voets T (2014) Peripheral thermosensation in mammals. Nat. Rev. Neurosci. 15: 573–589 Available at: http://dx.doi.org/10.1038/nrn3784

Wang D, O’Halloran D & Goodman MB (2013) GCY-8, PDE-2, and NCS-1 are critical elements of the cGMP-dependent thermotransduction cascade in the AFD neurons responsible for C. elegans thermotaxis. J. Gen. Physiol. 142: 437–49 Available at: http://www.jgp.org/lookup/doi/10.1085/jgp.201310959 [Accessed November 26, 2013]

Wasserman SM, Beverly M, Bell HW & Sengupta P (2011) Regulation of response properties and operating range of the AFD thermosensory neurons by cGMP signaling. Curr. Biol. 21: 353–362 Available at: http://www.pubmedcentral.nih.gov/articlerender.fcgi?artid=3057529&tool=pmcentrez&rendertype=abstract [Accessed October 8, 2014]

Yu Y V., Bell HW, Glauser DA, Hooser SD Van, Goodman MB, Sengupta P, VanHooser SD, Goodman MB & Sengupta P (2014) CaMKI - Dependent Regulation of Sensory Gene Expression Mediates Experience - Dependent Plasticity in the Operating Range of a Thermosensory Neuron. Neuron 84: 919–926 Available at: http://dx.doi.org/10.1016/j.neuron.2014.10.046

